# GenomicSEM Modelling of Diverse Executive Function GWAS Improves Gene Discovery

**DOI:** 10.1101/2024.02.12.579907

**Authors:** Lucas C Perry, CHARGE Consortium, Nicolas Chevalier, Michelle Luciano

## Abstract

Previous research has supported the use of latent variables as the gold-standard in measuring executive function. However, for logistical reasons genome-wide association studies (GWAS) of executive function have largely eschewed latent variables in favour of singular task measures. As low correlations have traditionally been found between individual executive function (EF) tests, it is unclear whether these GWAS have truly been measuring the same construct. In this study, we addressed this question by performing a factor analysis on summary statistics from eleven GWAS of EF taken from five studies, using GenomicSEM. Models demonstrated a bifactor structure consistent with previous research, with factors capturing common EF and working memory-specific variance. Furthermore, the GWAS performed on this model identified 20 new genomic risk loci for common EF and 4 for working memory reaching genome-wide significance beyond what was found in the constituent GWAS, together resulting in 29 newly mapped EF genes. These results help to clarify the underlying genetic structure of EF and support the idea that EF GWAS are capable of measuring genetic variance related to latent EF constructs even when not using factor scores. Furthermore, they demonstrate that GenomicSEM can combine GWAS with divergent and non-ideal measures of the same phenotype to improve statistical power.

## Introduction

Executive function (EF) covers the higher-order cognitive processes used to engage in goal-directed behaviour (Ahmed & Miller, 2011, Welsh et al., 2006). It has a role in promoting academic achievement (St Clair-Thompson & Gathercole, 2006; Best et al., 2011; Deer et al., 2020) and health behaviours (Reimann et al., 2020), and has been implicated across a wide variety of psychiatric diagnoses including ADHD (Brown, 2009; Marije Boonstra et al., 2005), schizophrenia (Johnson, Selfridge & Zalewski, 2001; Wobrock et al., 2009; Orellana & Slachevsky, 2013) and depression (Fossati et al., 2002; Letkiewicz et al., 2014; Cotrena et al., 2016). Multiple twin studies have suggested that EF is fully heritable at the latent variable level in adolescents (Friedman et al., 2008; Engelhardt et al., 2015), seemingly violating the supposed ‘law’ of behavioural genetics that no trait is 100% heritable (Plomin & Deary, 2015). Although twin studies do support an increased role for environment in predicting EF during adulthood (Friedman et al., 2016; Gustavson et al., 2018; Morrison et al., 2021), they still support additive genetic effects as the primary source of stability in EF over time. Furthermore, twin studies show that genes can explain much of the association between EF and psychiatric diagnoses (Harden et al., 2020), stressful life events (Morrison et al., 2021), and math and reading ability (Daucourt et al., 2020). Thus, understanding the relation between EF and outcomes of interest requires a strong understanding of the role that genes play in EF development.

A key piece of theory underlying our understanding of EF is the unity-diversity model, which is the most well-cited model of EF in the literature (Miyake et al., 2000). This model holds that EF is comprised of the three separable factors known as inhibition, updating, and shifting, which share common variance known as common EF. This model has been subsequently revised to a bifactor model, consisting of a common EF factor onto which all items load as well as orthogonal updating-specific and shifting-specific factors (Miyake & Friedman, 2012; Friedman & Miyake, 2017). Notably, this model lacks an inhibition specific factor, as the authors argue there is no inhibition-specific variance. This model has important implications for the measurement of EF, as it has been argued that on a theoretical level, an EF test measuring only EF is impossible. This is because EF is only engaged when directing cognitive resources towards specific tasks, which carry their own cognitive demands that are also captured in task performance (Miyake and Friedman, 2012). The need to eliminate this so-called ‘task impurity’, as well as measurement error, has led to the best-practice recommendation that EF be measured by multiple EF tests combined into a latent variable model like unity-diversity through CFA or similar methods (Friedman & Miyake, 2017). Importantly, latent EF factors typically outperform individual EF tests in predicting other outcomes of interest (Harden et al., 2020; Gustavson et al., 2022), and produce stronger correlations for EF assessments performed in different contexts (Freis et al., 2023), emphasizing the value of this measurement model. However, there is little-to-no agreement upon a gold-standard set of EF tasks for researchers to use, either when building these factor scores or as single-test measures. A literature review of empirical studies of EF published between 2008 and 2013 identified 109 tasks used for assessment throughout the field, of which 53 were used in multiple studies (Baggetta & Alexander, 2016). The use of such a diverse array of tests might not be an issue if they were highly correlated measures, however this is not the case. Perhaps owing to task impurity, the individual EF tests used in CFAs typically correlated below 0.4, with several showing no significant correlation (Lehto et al., 2003; Miyake et al., 2000). Despite these issues, single-task measurement of EF remains popular in clinical research, a methodological challenge meta-analyses often have to contend with (Andrews et al., 2021; Khoury et al., 2015; Lund et al., 2022; Power et al., 2021). Researchers cannot be sure that correlations with single-test EF measures are actually due to EF, nor that the results will replicate when performed with different EF measures, making the interpretation of any EF study using such measures challenging.

These issues are problematic for genome-wide association studies (GWAS) of EF, where the logistics of the large cohorts required for sufficient statistical power pressures researchers into using abbreviated test batteries that are quicker and easier to administer. As a consequence, marked heterogeneity exists in the approaches to measuring EF phenotypes used in GWAS. Across the 7 GWAS of EF as measured through cognitive tests that could be identified in the literature, only two, trail-marking part B and backwards digit span, were used in more than one study (Ibrahim-Verbaas et al., 2016; Zhang et al. 2018; Donati, Dumontheil, and Meaburn, 2019; Wendel et al., 2021; Hatoum et al, 2023; Dueker et al., 2023; Arnatkeviciute et al., 2023). As these GWAS have largely been performed on separate, weakly-correlated tests rather than latent factors, it is difficult to determine if they are measuring constructs similar to those found elsewhere in EF literature or even in other GWAS. One notable exception is a study conducted in the UK Biobank, which is both the largest GWAS of EF to date (N> 400,000) and the only such study to use a factor score (Hatoum et al, 2023). However, the UK Biobank cognitive test battery was not designed to measure EF, with their suitability to do so being based on a post-hoc analysis of their correlations with common reference EF tests (Fawns-Ritchie & Deary, 2020). The battery included only two traditional EF tasks, the trail-making task and a backwards digit span. As a consequence of these limitations, the factor analysis derived from the UK Biobank’s cognitive tests was only able to capture common EF, and the subsequent polygenic score was not a predictor of the updating or shifting-specific factors (Hatoum et al, 2023). And while the polygenic score was able to predict the common EF factor, its predictive power (β = 0.171, partial r = 0.136) (Hatoum et al., 2023) is still far below what twin studies of EF imply should be predictable from genetics (Friedman et al., 2008; Engelhardt et al., 2015), even considering the limitations of SNP heritability. But larger GWAS meta-analyses cannot be performed with measurement of the phenotype so fractured. Further, measurement error reduces statistical power in GWAS (Liao et al., 2014, van der Sluis et al., 2010), which may account for the failure of EF GWAS not using factor scores to find any significant associations. There is thus good reason to think measuring EF through latent factors will improve the quality of EF GWAS above and beyond what is possible simply by increasing sample size of single-test measures.

GenomicSEM provides a powerful tool to address both the need for latent variables and larger sample sizes in EF GWAS. This method allows for the analysis of multiple GWAS using summary statistics, using them to build a genetic covariance matrix using Linkage Disequilibrium-score regression (LDSC) (Bulik-Sullivan et al., 2015), analogous to the covariance matrix employed by traditional structural equation modelling (Grotzinger et al., 2019). Therefore, rather than requiring building a CFA from multiple EF tests administered to the same participants, GenomicSEM allows a CFA to be built from GWAS of single-test measures without requiring participant overlap. Furthermore, once a model is accepted, GenomicSEM enables users to run a multivariate GWAS on the specified latent factors, using only the summary statistics used to build the model rather than individual-level genetic data. This is accomplished by fitting a structural equation model for each SNP in the analysis, using the SNP effects included in the genetic and sampling covariance matrices to estimate the influence of each SNP on the latent factors (Grotzinger et al., 2019). Thus, EF GWAS ostensibly measuring the same EF component but unsuitable for meta-analysis due to their use of different measures can be combined at the latent factor level, providing an alternative path for increasing sample size while also addressing task impurity. GenomicSEM does not model shared variance due to environment, which is an important limitation of the method compared to traditional SEM. However, this also provides a method of addressing concerns about correlations between tests representing a form of measurement error (Willoughby et al., 2014; Willoughby et al., 2016; Camerota et al., 2020), which should not appear at the genetic level across distinct samples.

The present study represents the first effort to model multiple latent EF factors in GenomicSEM, as well as the first EF model to use GWAS summary statistics taken from different samples. The purpose of the present study will therefore be to examine the factor structure of the currently available EF GWAS, determining the extent to which EF measures from separates studies load onto common constructs. Furthermore, where such common constructs are found, we aim to use GenomicSEM to perform GWAS on the latent factors, to increase statistical power for gene discovery beyond that of the constituent GWAS and allowing for the creation of polygenetic scores of EF factors that should be more predictive of latent EF factors than those derived from the individual studies.

## Method

### Summary Statistics

A literature search was conducted in order to identify GWAS of EF suitable for inclusion in GenomicSEM. Inclusion criteria were EF measured by a cognitive test that showed statistically significant SNP heritability in LDSC (Bulik-Sullivan et al., 2015). As ethnically diverse samples are currently incompatible with GenomicSEM (Grotzinger et al., 2019), samples also needed to be either conducted in a solely European sample or have a European subsample available to meet inclusion criteria. Other demographic features such as age were not part of inclusion criteria. The summary statistics which met inclusion criteria are presented in Table 1. While summary statistics for the Wisconsin Card Sorting Test (Zhang et al. 2018), the PsyCourse Study’s Trail Making Part B and Backwards Digit Span, (Wendel et al., 2021) and the CHARGE consortium’s Trail-Making Part B were also obtained, these failed to show significant SNP heritability in LDSC and so were excluded from further analysis. In addition to their factor score GWAS statistics, Hatoum et al. (2023) made available summary statistics for GWAS of the individual tests that contributed to it. As these tests should in theory tap different EF domains, we used these separate summary statistics when building our model. RsIDs for Arnatkeviciute et al. (2023) were obtained using bedops (Neph et al., 2012). Results for the UK Biobank’s Symbol Digit Substitution Test, the CHARGE consortium’s Stroop test, the Stop-Signal test, and both principal components form the NIHR Bioresource were reverse-coded so all positive loadings would correspond to better task performance.

**Table 1:**
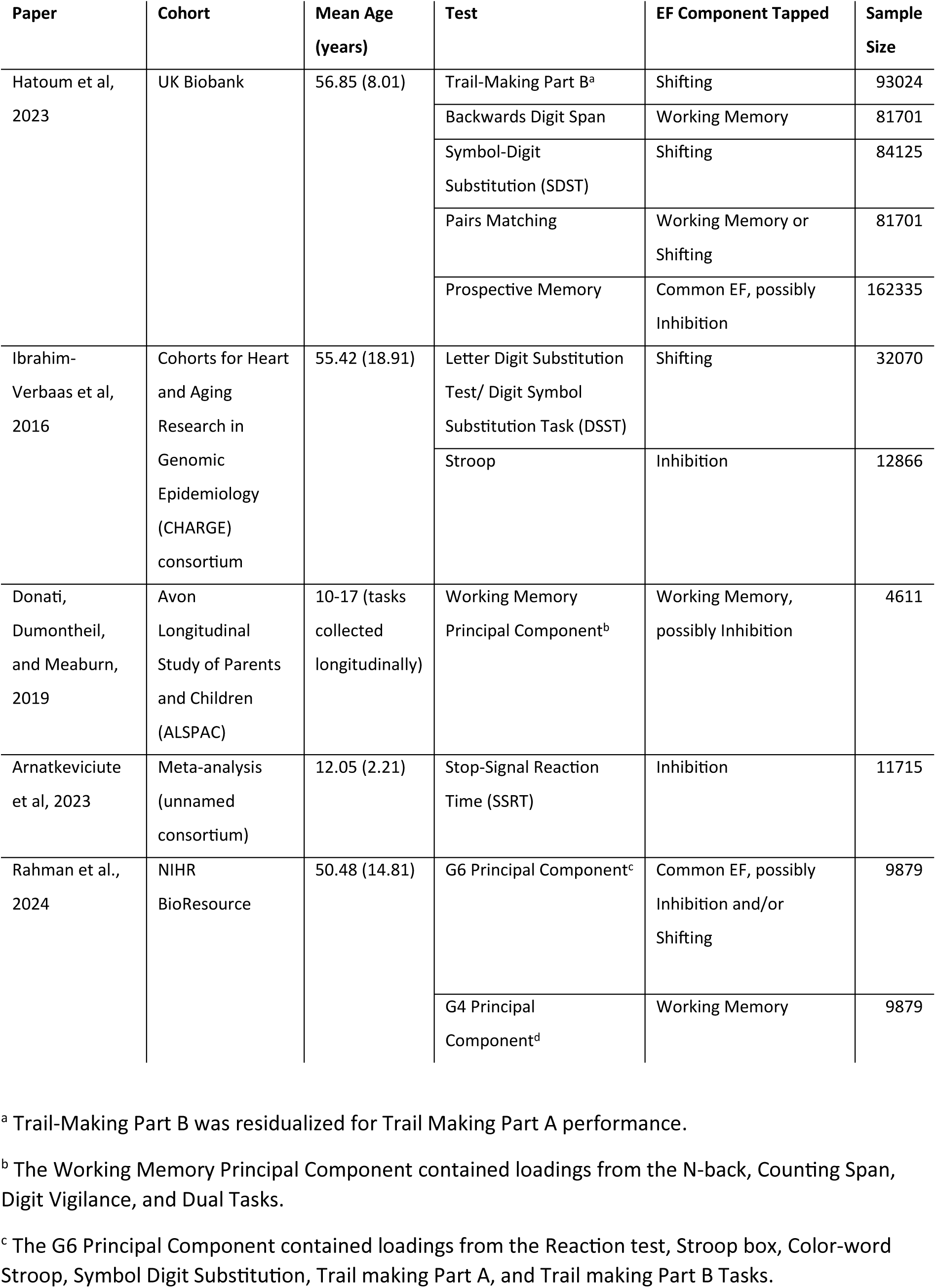
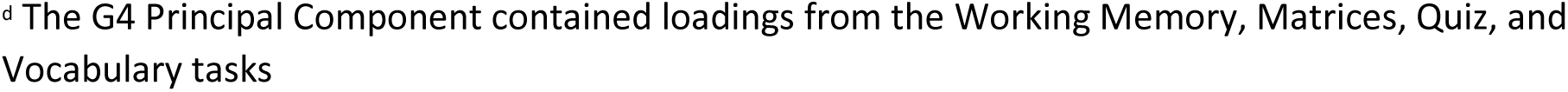
GWAS summary statistics for EF test variables used in the present study.

| Paper | Cohort | Mean Age (years) | Test | EF Component Tapped | Sample Size |
| --- | --- | --- | --- | --- | --- |
| Hatoum et al, 2023 | UK Biobank | 56.85 (8.01) | Trail-Making Part B<sup>a</sup> | Shifting | 93024 |
|  |  |  | Backwards Digit Span | Working Memory | 81701 |
|  |  |  | Symbol-Digit Substitution (SDST) | Shifting | 84125 |
|  |  |  | Pairs Matching | Working Memory or Shifting | 81701 |
|  |  |  | Prospective Memory | Common EF, possibly Inhibition | 162335 |
| Ibrahim-Verbaas et al, 2016 | Cohorts for Heart and Aging Research in Genomic Epidemiology (CHARGE) consortium | 55.42 (18.91) | Letter Digit Substitution Test/ Digit Symbol Substitution Task (DSST) | Shifting | 32070 |
|  |  |  | Stroop | Inhibition | 12866 |
| Donati, Dumontheil, and Meaburn, 2019 | Avon Longitudinal Study of Parents and Children (ALSPAC) | 10-17 (tasks collected longitudinally) | Working Memory Principal Component<sup>b</sup> | Working Memory, possibly Inhibition | 4611 |
| Arnatkeviciute et al, 2023 | Meta-analysis (unnamed consortium) | 12.05 (2.21) | Stop-Signal Reaction Time (SSRT) | Inhibition | 11715 |
| Rahman et al, 2024 | NIHR BioResource | 50.48 (14.81) | G6 Principal Component<sup>c</sup> | Common EF, possibly Inhibition and/or Shifting | 9879 |
|  |  |  | G4 Principal Component<sup>d</sup> | Working Memory | 9879 |

<sup>a</sup> Trail-Making Part B was residualized for Trail Making Part A performance.
<sup>b</sup> The Working Memory Principal Component contained loadings from the N-back, Counting Span, Digit Vigilance, and Dual Tasks.
<sup>c</sup> The G6 Principal Component contained loadings from the Reaction test, Stroop box, Color-word Stroop, Symbol Digit Substitution, Trail making Part A, and Trail making Part B Tasks.
<sup>d</sup> The G4 Principal Component contained loadings from the Working Memory, Matrices, Quiz, and Vocabulary tasks

### Factor structure

While it was expected that all terms would capture common EF, we additionally assigned each set of GWAS summary statistics loadings onto one of three EF factors-inhibition, working memory, or shifting. Available data were insufficient to distinguish updating from other forms of working memory, so a general working memory factor took the place of updating in our model. While many of the tests presented in Table 1 have long been recognized in the literature as tapping specific EF subcomponents, Hatoum et al.’s (2023) GWAS of the UK Biobank included nonstandard tests for which their capacity to measure EF was established based on correlations with standard EF reference tests (Fawns-Ritchie & Deary, 2020). As the reference tests used tap different EF subcomponents (shifting for Trail Making Part B, working memory for Backwards Digit Span, and complex EF for Tower Test) we used a combination of the relative difference in correlations with these tests as reported in that paper and the demands apparent from the tasks’ design to assign the nonstandard UK Biobank tests to specific EF domains.

Symbol-Digit substitution correlated strongly with the Trails reference tests (r = –0.558, p < .001) and only weakly with Backwards Digit Span (r = 0.177, p < .05), indicating that it may contain significant shifting-specific variance (Fawns-Ritchie & Deary, 2020). The Charge consortium also included a similar GWAS combining Letter Digit Substitution and Digit Symbol Substitution, although they used it as a processing speed test (Ibrahim-Verbaas et al, 2016). While this is a common interpretation in the literature, some studies have also supported a relationship between substitution tasks and set shifting, including when measured through Trails B (Jehu et al., 2021; Knowles et al., 2015; Lessov-Schlaggar et al., 2007), which may reflect the need to switch between mental sets of symbol-digit pairings across trials. As such, while we assigned both digit substitution GWAS to set shifting, we also included an orthogonal factor which loaded these GWAS alone, in order to minimize variance due to processing speed captured by our shifting factor.

Pairs matching correlated modestly with Trails B (r = 0.277, p < .001) but had no significant correlation with Backwards Digit span (r = –0.122) suggesting a weaker but still shifting-specific EF component (Fawns-Ritchie & Deary, 2020). This is however surprising, as the task, which involves the participant memorizing an arrangement of cards and then selecting match pairs after they have been flipped over, has apparent working memory demands but no clear link to shifting. As such, we tested models assigning it to both working memory and shifting.

Prospective memory had modest correlations with all EF reference tests (Backwards Digit Span r = 0.265 p < .01, Trails B –0.306, p < .001) consistent with literature finding prospective memory to be associated with multiple EF domains (Kerns, 2000; Shum et al., 2008; Yang et al., 2011). However, we theorized that the task would be most strongly related to inhibition, which was not represented in the reference tests. The UK Biobank’s prospective memory task is event-based, a classification which previous research has suggested is most strongly related to inhibition (Mahy et al., 2014; Schnitzspahn et al., 2013; Yi et al., 2014; Zuber et al., 2019). Furthermore, the task is conceptually similar to inhibition tasks such as stop-signal and Go/No-Go, both of which are reaction tasks in which a response is inhibited upon receiving a certain signal (Raud et al., 2020). For the prospective memory task, the reaction is to simple onscreen instructions (e.g., ‘touch the blue square’), with the inhibition signal being a previously given set of countermanding instructions (e.g., ‘When you are told to touch the blue square, touch the orange circle’). While the perception of the signal is dependent upon the participant’s prospective memory, responding appropriately to it nevertheless has clear inhibitory demands (Hatoum et al, 2023).

The working memory principal component extracted from the ALSPAC cohort (Donati et al. 2019), while in some ways similar to a CFA approach, comes with issues which require a more nuanced interpretation. PCA is not hypothesis-driven, and analyses all variance in the terms rather than only their shared variance. Consequently, the working memory principal component did not only load onto tasks with obvious working memory demands, but instead had loadings on the Digit Vigilance and Dual Tasks, which are respectively tasks of sustained and divided attention (Donati et al., 2019). In an earlier version of their analysis, the authors also found loadings from the stop-signal task variables at both age 10 and 15 onto their working memory principal component (Donati et al., 2019). While these variables were removed due to their loading onto multiple components, they nevertheless indicated that the principal component they identified as working memory shared variance with the stop-signal task, possibly indicating that it captured variance due to inhibition or some other shared cognitive process. The authors of that paper offered a similar explanation for the principal component they identified as inhibition showing no SNP heritability, suggesting that this may be due to there being no genetic variance unique to inhibition (Donati et al., 2019). Perhaps as a result of these features, during our preliminary analysis with the Stop-Signal Task included, the Stop-Signal, ALSPAC working memory component, and prospective memory showed a relationship which was inconsistent with inhibition or common EF (see supplemental materials). As such, when the decision was made to remove the Stop-Signal, we hypothesized that ALSPAC working memory and prospective memory would retain some relationship. We accounted for this through a correlated error term, declining to assign an interpretation without sufficient conceptual clarity, and retained it in the model as it was found to improve fit.

Finally, the NIHR Bioresource study also included two principal components with potential EF demands (Rahman et al., 2024). While these were conceptualised by the paper as measures of *g*, their use of rotation to produce two factors is outside the norm of *g* research, where *g* is typically defined as the first unrotated principal component of a battery of cognitive tests (Plomin & Deary, 2015). As these components included loadings from common EF tests, we hypothesized that they may have captured EF variance as well. G6 contained loadings with two types of Stroop, Trails, as well as Symbol Digit Substitution. As such, we hypothesized that this term would contain common EF demands as well as possible inhibition or shifting-specific demands, although due to its multicollinearity with several other terms in LDSC it could not be included in our tested models. G4 contained loadings from vocabulary and fluid intelligence tasks more in line with *g* research, but also a loading from a digit span task they identified as working or numerical memory. As such, we hypothesized that the term would have working memory demands. However, it should be acknowledged that said working memory task had the weakest loading of the four terms comprising G4 (∼0.17) and consequentially a phenotypic correlation of 0.57 with the G4 principal component. As such, we compared our accepted model to one which dropped the G4 term entirely in order to ensure that the term aligned in its demands with the rest of the model.

### Analysis

Analysis was conducted in GenomicSEM (Grotzinger et al., 2019). Models were tested using the absolute fit thresholds proposed in the original GenomicSEM paper, with CFI values of ≥0.95 considered good fit and ≥0.90 considered acceptable fit (Kenny, 2014), and values of SRMR of <0.05 considered good fit and <0.10 considered acceptable fit (Hoyle, 1995). The genetic covariance matrix was obtained using LDSC (Bulik-Sullivan et al., 2015); genetic correlations are presented in Table 2. Our initial hypothesis was that the data would fit a unity-diversity three-factor model of set shifting, inhibition, and working memory. We tested both the standard version of this model and a version including an orthogonal factor for substitution task –specific variance. We also tested all possible combinations of models merging two EF factors together. Finally, bifactor models, which added an orthogonal common EF factor to the previously described models, were also tested, both including or eliminating inhibition-specific variance.

**Table 2:**
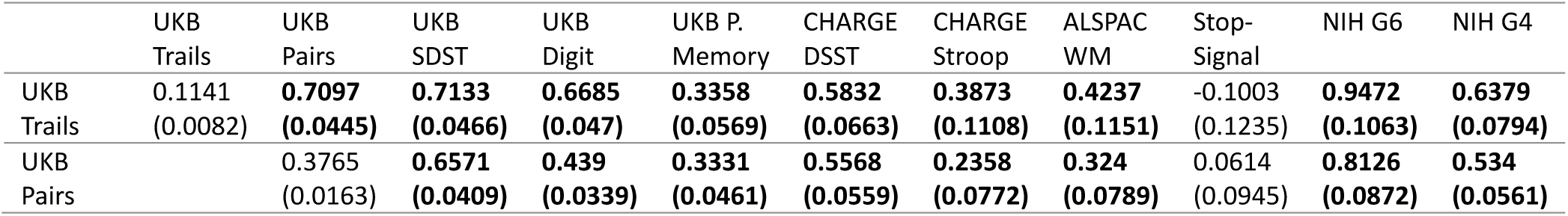
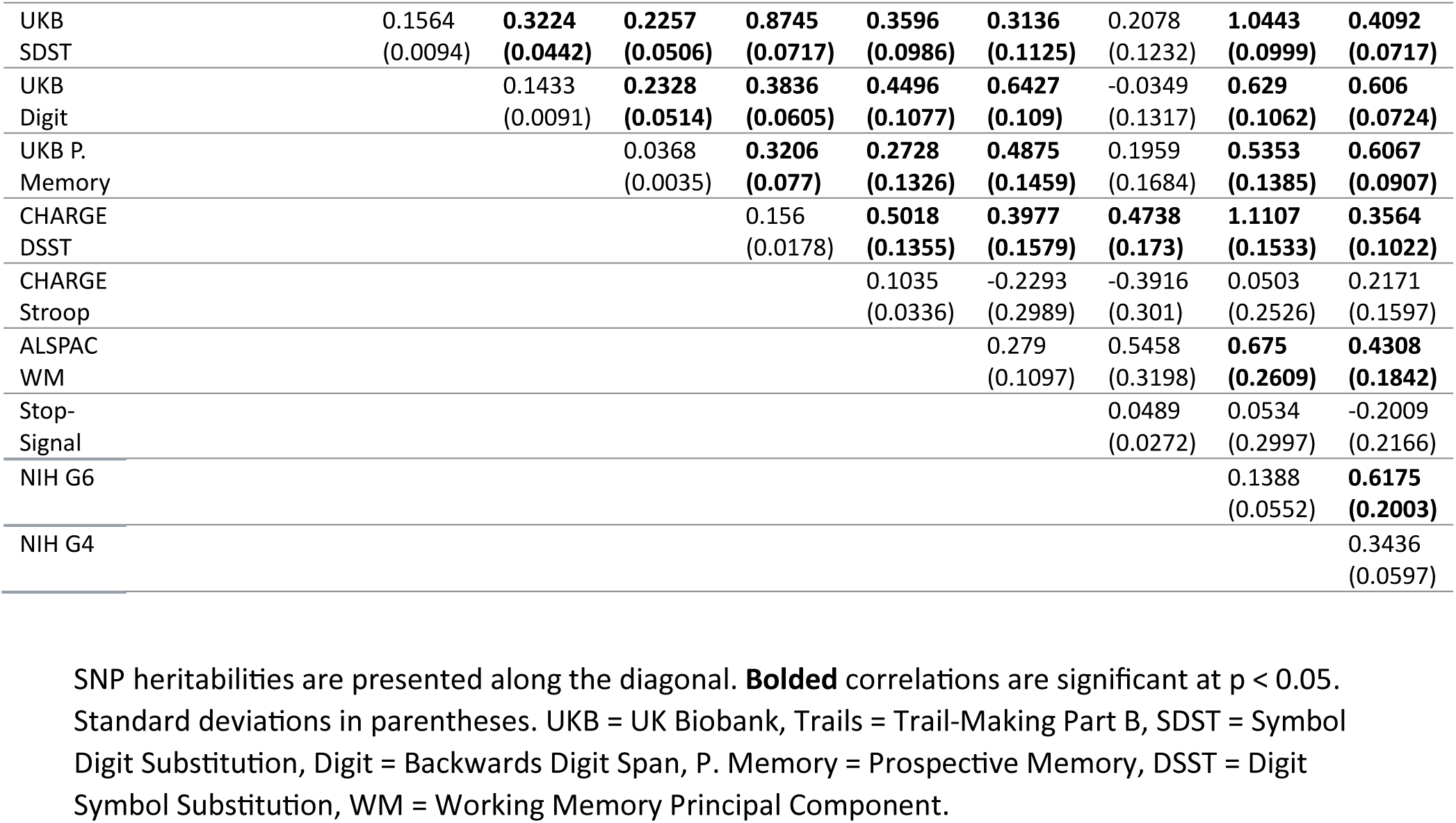
Genetic correlation between GWAS summary statistics.

The accepted model was run through the user-specified GWAS function in GenomicSEM, regressing SNPS onto each of the latent factors. The genetic covariance matrix was smoothed due to significant differences in *n* between samples, and SNPs which showed a difference of more than 1 in Z-score between the pre– and post-smoothed genetic covariance matrices were removed from the results. SNPs that showed a Q-SNP heterogeneity P-value < 5 x 10^-08^ were also removed.

LDSC (Bulik-Sullivan et al., 2015) was used to obtain intercepts and genetic covariance with other psychiatric phenotypes, including Hatoum et al.’s (2023) full-sample EF and processing speed (taken from the UK Biobank’s snap game) summary statistics, as well as summary statistics for educational attainment (Okbay et al., 2022), cognitive performance (Lee et al., 2018), and 9 different disorders taken from the psychiatric genetics consortium (Arnold et al., 2018; Demontis et al., 2023; Forstner et al., 2021; Grove et al., 2019; Howard et al., 2019; Mullins et al., 2021; Trubetskoy et al., 2022; Watson et al., 2019; Yu et al., 2019). In line with the approach taken by Hatoum et al. (2023) variance shared between our EF factors, general cognitive ability (UK Biobank’s verbal-numerical reasoning test), and processing speed (snap game) was controlled for using multiple regression in genomicSEM, allowing us to examine the correlations between variance unique to our factors and outcomes of interest. GWAS results were entered into FUMA/MAGMA (Watanabe et al., 2017) to obtain functional annotations and gene mappings. These results were compared to the publicly available FUMA/MAGMA results from Hatoum et al.’s (2023) full sample and the two constituent GWAS which produced significant results (CHARGE DSST and NIH G4) so that overlaps could be removed and a list of novel associations obtained. To filter lead and independent significant SNPS, we first removed all significant SNPs with rsIDs matching significant SNPs in the previous EF GWAS, then used the LD files outputted by FUMA to identify and remove any significant SNPs in our GWAS which were in LD (*r^2^* > 0.6) with significant SNPs from the previous EF GWAS. For gene mappings, we likewise removed any genes with gene symbols matching previous EF associations, as well as any genes associated with the previously removed SNPs. For genomic risk loci, we used the GenomicRanges package in R to identity and remove risk loci overlapping those found in previous EF GWAS (Lawrence et al., 2013).

## Results

Table 2 presents genetic correlations for all summary statistics which met inclusion criteria by showing significant SNP heritability in LDSC. With the exception of Stop-Signal, all summary statistics showed multiple significant genetic overlaps with other measures of EF, indicating that they tap a common phenotype. However, the NIHR G6 displayed multicollinearity with two other terms, and was consequentially removed from the analysis.

### Confirmatory factor analysis

Fit statistics for our CFAs are presented in Table 3. During modelling, the Stop-Signal Reaction Time measure failed to significantly load onto Common EF in bifactor formulations, or inhibition in correlated-factors formulations (see supplemental material). As it also did not significantly correlate with any other measure except for the CHARGE consortium’s DSST, it is excluded from all models presented here.

**Table 3:** Confirmatory factor analysis model results for the genomic structure of EF.

| **Model** | **$\chi^2$** | **df** | **$\chi^2$ p-value** | **AIC** | **CFI** | **SRMR** |
| --- | --- | --- | --- | --- | --- | --- |
| Common Factor Model | 236.1202 | 27 | 2.776111e-35 | 272.1202 | 0.895631 | 0.1306878 |
| Three Factor correlated | 219.4508 | 23 | 5.467484e-34 | 263.4508 | 0.9019541 | 0.1246057 |
| Three factor correlated with Substitution | 158.1513 | 20 | 1.706908e-23 | 208.1513 | 0.9310506 | 0.1131072 |
| Bifactor with Working Memory | 186.1265 | 22 | 5.753337e-28 | 232.1265 | 0.9180867 | 0.1081619 |
| *Bifactor With Working Memory and Substitution* | *47.0705* | *19* | *0.0003489703* | *99.0705* | *0.9859904* | *0.09571648* |
| Bifactor with WM, Substitution is Shifting | 47.11906 | 19 | 0.0003434276 | 99.11906 | 0.9859662 | 0.1049589 |
| Bifactor drop Pairs from WM | 75.56783 | 20 | 2.190229e-08 | 125.5678 | 0.9722669 | 0.09663167 |
| Bifactor, Prospective loads onto WM | 64.11958 | 20 | 1.610782e-06 | 114.1196 | 0.9779805 | 0.104505 |
| Bifactor with WM, Substitution is a correlated residual | 129.9266 | 21 | 1.047574e-17 | 177.9266 | 0.9456362 | 0.09716726 |

Endorsed model in italics.

The basic three-factor model met acceptable fit thresholds by CFI but failed to do so for SRMR (see Figure 1). Adding the orthogonal substitution-specific factor slightly improved fit, although it remained below acceptable thresholds (this model also required the residual variance for the CHARGE DSST to be fixed to 0 to avoid empirical under-identification, as no EF terms correlated with the substitution factor). Fit was likewise improved slightly by the bifactor model with working memory, although this, too, fell below acceptable thresholds. After the addition of the orthogonal substitution-specific factor (see Figure 2), the resulting model fit improved significantly, resulting in good fit by CFI and acceptable fit by SRMR, as well as a statistically significant improvement over both the common factor and previous bifactor model by χ2 and AIC.

**Figure 1:**
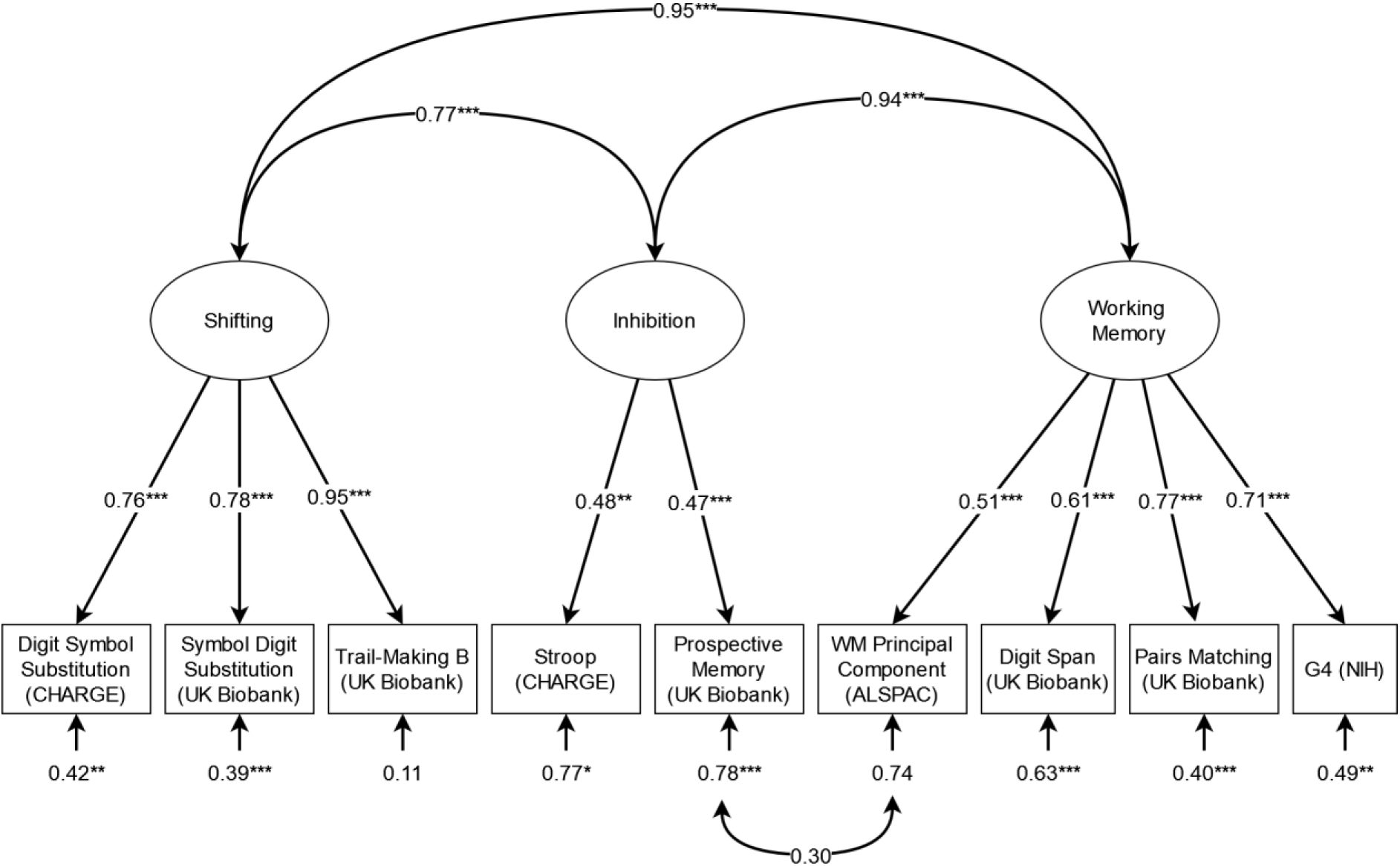
Standardised results for the initial three-factor confirmatory factor analysis model of EF GWAS. *p < .05, ** p < .01, *** p < .001.

**Figure 2:**
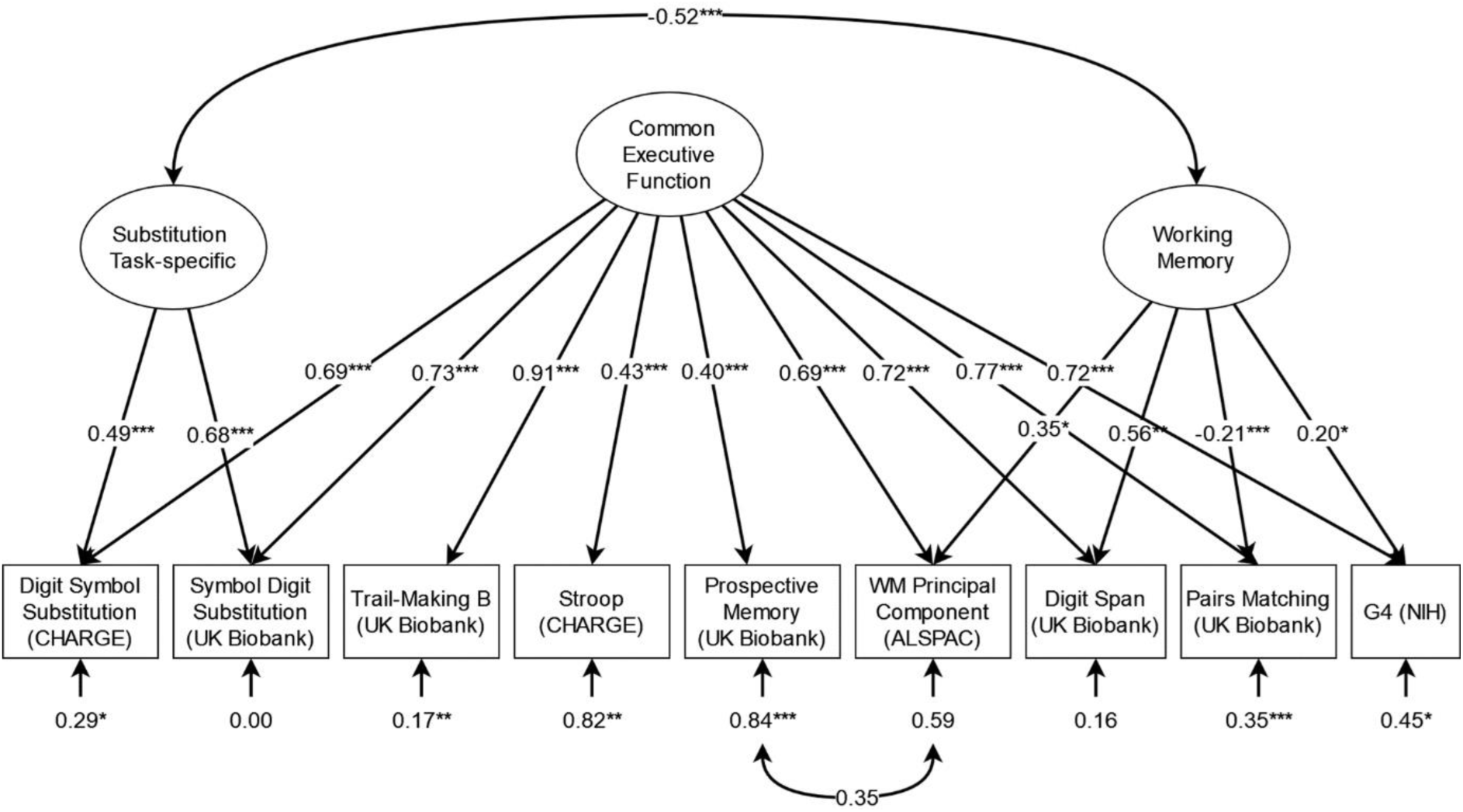
Standardised results for the accepted confirmatory factor analysis model of EF GWAS. *p < .05, ** p < .01, *** p < .001.

Using this adequate-fitting model as a basis, we tested several modifications to confirm our model decisions. We tested the possibility that the substitution factor might instead be characterized as shifting with a model loading Trail Making onto said factor as well. However, while model fit was similar, we found Trail Making loaded only weakly (∼0.16) and nonsignificantly onto the substitution factor, making its identification with shifting dubious and giving us clear reason to prefer the more parsimonious model without this loading. Given that Pairs matching unexpectedly correlated negatively with working memory, we tested the possibility that it contained only common EF or shifting variance by dropping it from working memory, but found this worsened fit by χ2 and AIC. We also tested a model where prospective memory was assigned to the working memory factor instead of being given a correlated residual with ALSPAC Working Memory, but found this, too, worsened fit. Finally, we tested a model which attempted to account for the substitution factor with a correlated residual, but found that while it was an improvement in fit compared to the bifactor model without the substitution factor, it was still a worse fit than the model with that factor. As none of our modifications were found to improve fit, the bifactor model with working memory and substitution-specific variance was accepted and used for the GWAS.

As some question remained about the inclusion of the NIH G4 principal component in our model, we fit the accepted model without that term which produced similar fit (CFI = 0.995, SRMR = 0.090). As the resulting summary statistics of the GWAS of the models with and without NIH G4 were at near-to-complete unity in LDSC (0.993 ±0.048 for common EF and 1.003 ±0.083 for working memory) we chose to focus on the more powerful GWAS including the NIH G4 principal component.

### GWAS

Table 4 presents a brief summary of our GWAS results after annotation in FUMA. We also present a summary of the subset of our results which were not found in the constituent EF GWAS. Full FUMA and MAGMA results are available in supplementary tables S1 –S20, and include lead and independent significant SNPS, genomic risk loci, identified genes and GWAS Catalog mappings, both with and without overlaps with previous EF GWAS.

**Table 4:** Summary of GWAS Results for the Common Factor Model, with and without overlaps with previous EF GWAS.

| GWAS | Genomic Risk Loci | Lead SNPs | Independent Significant SNPs | Mapped Genes | Top Gene |
| --- | --- | --- | --- | --- | --- |
| Common EF | 46 | 47 | 101 | 163 | *EXOC4* |
| Novel EF | 20 | 15 | 27 | 22 | *MON1A* |
| Working Memory | 5 | 5 | 5 | 12 | *LRP8* |
| Novel WM | 4 | 4 | 4 | 7 | *LRP8* |

### LDSC

LDSC results for the common EF and working memory-specific factors are presented in Table 5. We also present genetic association results after adjusting for overlap with GWAS of general cognitive ability and processing speed via multiple regression, using summary statistics from Hatoum et al. (2023).

**Table 5:** Genetic correlations between executive function and other cognitive phenotypes.

|  | Common EF | Working Memory | Common EF Regressed | Working Memory Regressed |
| --- | --- | --- | --- | --- |
| cEF (UK Biobank) | **0.9645**<br>(0.0462) | **0.3333**<br>(0.0369) |  |  |
| General Cognitive Ability | **0.8572**<br>(0.0358) | **0.6315**<br>(0.0428) |  |  |
| Processing Speed | **0.2459**<br>(0.0277) | 0.0471<br>(0.0357) |  |  |
| Educational Attainment | **0.4733**<br>(0.0236) | **0.4712**<br>(0.0309) | **-0.3846**<br>(0.1014) | 0.0659<br>(0.0406) |
| ADHD | **-0.2689**<br>(0.0302) | **-0.2093**<br>(0.0445) | 0.1820<br>(0.1472) | 0.0849<br>(0.0933) |
| Autism | 0.0832<br>(0.0382) | **0.171**<br>(0.0509) | -0.3138<br>(0.1796) | 0.0654<br>(0.0714) |
| Schizophrenia | **-0.2788**<br>(0.0271) | 0.0338<br>(0.0357) | **-0.2202**<br>(0.1017) | **0.2876**<br>(0.0692) |
| Bipolar Disorder | **-0.2036**<br>(0.0289) | 0.0908<br>(0.0428) | **-0.4215**<br>(0.1302) | 0.2348<br>(0.0778) |
| Anxiety | *-0.188*<br>(0.0691) | 0.1396<br>(0.0931) | -0.4681<br>(0.2521) | *0.2959*<br>(0.1476) |
| Anorexia Nervosa | 0.0528<br>(0.0426) | **-0.2927**<br>(0.0571) | *0.4966*<br>(0.1918) | **-0.3808**<br>(0.1028) |
| Major Depressive Disorder | **-0.1464**<br>(0.0253) | -0.011<br>(0.035) | -0.2688<br>(0.1259) | 0.0744<br>(0.0640) |
| Obsessive-Compulsive Disorder | -0.0781<br>(0.0622) | *0.2096*<br>(0.0891) | -0.3555<br>(0.2581) | *0.3073*<br>(0.1542) |
| Tourette Syndrome | 0.0027<br>(0.0458) | *0.1694*<br>(0.0629) | 0.1236<br>(0.2081) | *0.3371*<br>(0.1129) |

Regress columns control for overlap with cognitive ability and processing speed. Bolded results are statistically significant after adjusting for multiple testing (Bonferroni’s corrected p < 0.00192). Italicized results are nominally significant at p < 0.05. Standard deviations in parentheses.

## Discussion

The purpose of this study was to perform a confirmatory factor analysis on currently available summary statistics from GWAS of executive function, in order to determine if these studies have been measuring the same construct and allowing us to perform more powerful GWAS on the basis of these shared latent factors. Our analysis produced a bifactor factor model largely in line with previous research, with orthogonal factors for common EF and working-memory-specific variance. That these results could be obtained on the basis of genetic correlations between tests performed largely on unrelated individuals suggests that the unity-diversity model of EF likely captures real biological processes that are reflected in performance on EF tests.

However, our results are unusual with respect to the Stop-Signal test (Arnatkevicuite et al, 2023) which we found had no significant genetic relationship to any other EF test except for the CHARGE consortium’s DSST, and no significant loading onto common EF. While Stop-Signal is a popular measure of inhibitory control, some EF CFAs have reported modest loadings for it (∼0.15), particularly for adolescent samples (Engelhardt et al., 2015; Freis et al., 2022). And while this may be sufficient to explain the term’s non-significant loading onto common EF, the negative relationship in LDSC between Stop-Signal and Stroop, another popular measure of inhibitory control, might suggest genetic separability beyond the low phenotypic loading. Again, the consortium sample used for the Stop-Signal GWAS skewed towards adolescents, in contrast to the older adult samples used for the UK Biobank and CHARGE, which raises the possibility that the genetic contribution to performance of this task may not be stable across age. Furthermore, if the previous correlation between Stop-Signal and other measures of inhibitory control have been primarily driven by the environment, Genomic SEM, which only takes genetic covariance into account, might present a very different relationship. Friedman et al.’s (2016) results in particular are consistent with this interpretation of our findings, as they found reduced loading for the Stop-Signal task onto inhibition at age 23, along with a loss of any statistically significant role for genetics in predicting its variance. Further research will be needed to disentangle why Stop-Signal produced these divergent results, and what implications this might hold for the task’s use in EF research more broadly.

### GWAS

We observed significant results for both common EF and working memory. As expected, given its large contribution to the sample, gene mappings for common EF showed a great deal of unity with the UK Biobank GWAS, including a shared top gene, *EXOC4.* Nevertheless, there were still 22 genes newly associated with EF for this factor, found neither in the UK Biobank nor in any other contributing GWAS. Furthermore, of the 27 novel independent significant SNPs for common EF, 15 were suggestively significant (p < 1 * 10^-6^) in Hatoum et al (2023). This demonstrates that the inclusion of these outside samples provided increased statistical power and new information, despite contributing only 59,426 additional participants to the GWAS. The top gene of the novel subset was *MON1A,* a gene believed to be involved in protein secretion that has been implicated in a number of other cognitive traits, including occupational attainment (Ko et al., 2022), household income (Hill et al., 2019), intelligence (Hill et al., 2019), creativity (Kim et al., 2024), and schizophrenia (Lam et al., 2019). In FUMA, the common factor also showed significant overlaps with gene-sets derived from GWAS of a number of cognitive phenotypes, most prominently brain morphology, autism, and schizophrenia. Working memory also produced GWAS hits, mapping to 12 genes including 7 not associated with constituent EF GWAS results. The top gene, *LRP8*, has been previously been associated with educational attainment (Lee et al., 2018; Okbay et al., 2022) and various changes in brain morphology (Fan et al., 2022; Naqvi et al., 2021; van der Meer et al., 2021), but interestingly not with any measure of general cognitive ability, suggesting a working memory-specific relationship with these changes may be more proximal.

### LDSC

Our common EF factor was at near-unity with the UK Biobank results in LDSC, showing that they measure the same phenotype. Consequentially, genetic associations are nearly identical to those reported by Hatoum et al (2023), with differences largely the result of the larger psychiatric genetics consortium studies that have since been made available. However, one noticeable difference is the increase in overlap with cognitive ability from 0.743 (0.013) to 0.857 (0.036). This may be a consequence of the inclusion of the NIH G4 principal component, which was conceived as a measure of *g* (although this term also produced a significant residual in our model, consistent with a partial separability of *g* and common EF). Despite this, associations after controlling for variation shared with cognitive ability and processing speed were of similar magnitude and direction to the UK biobank, although only associations with educational attainment and bipolar disorder survived correction for multiple testing. Associations with bipolar disorder were increased, suggesting that the genetic relationship between common EF and bipolar disorder is mediated by EF-specific genes. However, the association between common EF and educational attainment reversed and became negative. While similar results were reported by Hatoum et al (2023), the reasons for this finding are unclear. Previous research has suggested a trade-off between common EF and shifting-specific variance, resulting in better shifting being associated with worse outcomes in behavioural domains (Friedman et al., 2011; Herd et al., 2014), but to our knowledge there are no equivalent findings for common EF itself. These findings thus underscore the potential value of research into unique genetic contributions to cognitive phenotypes.

Our working memory factor showed a relatively modest association with the UK Biobank’s common EF but a stronger association with general cognitive ability, consistent with previous findings suggesting that working memory has a relationship to cognitive ability independent of common EF (Friedman et al., 2006). Said relationship appeared to drive the association with educational attainment, ADHD and autism, none of which remained after adjusting for overlap with cognitive ability and processing speed. However, working memory’s significant association with anorexia nervosa increased after the same adjustment, and it gained a significant association with schizophrenia as well as nominally significant associations bipolar disorder and anxiety. Notably, the direction of the effect for these new associations was opposite to that of the common EF, a surprising and difficult-to-interpret finding given that working memory deficits in these disorders are well established (Johnson’Selfridge & Zalewski, 2001; Soraggi-Frez et al., 2017; Saldarini et al., 2022). Nevertheless, these results suggest that variance shared with EF plays an important role in these disorders both via the common factor and through factor-specific variance.

### Limitations and Future Directions

Given the potential complexity implied by competing theories of EF, our model contained relatively few terms, constrained by the availability of appropriate GWAS. Our analysis was incapable of separating working memory into updating and maintenance factors, as some EF models do (Engelhardt et al., 2015; Wongupparaj et al., 2015), nor could it separate phonological from visuospatial working memory, which have been shown to have divergent relationships with several psychiatric diagnoses (Perugini et al., 2024; Zilles et al., 2012). One of the more unusual features of our model, the negative loading of pairs matching onto working memory, may be explainable by this distinction, as Pairs is the only visuospatial working memory task in the model. Changes in in the direction of the relationship between cognitive abilities when a common factor is extracted have been demonstrated in previous work (Knyspel & Plomin, 2024). As our bifactor model also extracted common variance (onto which Pairs loaded positively as expected), this may suggest trade-offs between the specific genetic components of visuospatial and phonological working memory. Future GWAS including a range of visuospatial working memory tasks will be important to enable these distinctions to be examined with more clarity.

The loss of Stop-Signal from the model limited our ability to specify an inhibition factor clearly in line with previous research, much less examine any of the proposed subdivisions (Friedman & Miyake, 2004; Stahl et al., 2014; Bender et al., 2016; Rey-Mermet et al., 2018). Future GWAS of additional inhibition and shifting tasks may enable the identification of genetic variance unique to these factors. Furthermore, our interpretation of the factors we were able to specify must also be treated with caution. Our summary statistics included GWAS of standard EF tests, nonstandard tests whose EF demands were established post-hoc, and principal components combining both EF and non-EF tests. Although they represent the best GWAS summary statistics currently available for measuring EF, it is nevertheless possible that the latent factors they tap may differ from those found in phenotypic literature. In considering these limitations and how they might be addressed in the future, we in fact highlight a major advantage of CFAs built in GenomicSEM over traditional CFAs.

Because the covariance matrix is built on the basis of genetic covariance rather than within-person covariance, terms from future GWAS of EF can be introduced as they become available, collaboratively iterating upon these results and in doing so increasing conceptual clarity.

A final limitation worth noting is the inclusion of GWAS only of European populations in our analysis. This limitation is unfortunately built into GenomicSEM itself, which requires that only GWAS of ethnically homogeneous samples be used. Two of the studies, which contributed GWAS summary statistics to the present study, Ibrahim-Verbaas et al, 2016 and Arnatkeviciute et al, 2023, made a point to construct their GWAS with multi-racial samples. This should be commended, as GWAS have historically and continue to overrepresent European populations in their sample (Popejoy & Fullerton, 2016; Rosenberg et al., 2010). However, because of the limitations of GenomicSEM, we were forced to use only the European subsample of these studies. Members of the GenomicSEM development team have indicated that GenomicSEM will in the future be improved to allow the inclusion of multiple LD files, making trans-ancestry GenomicSEM studies possible (*Multi-Ancestry LDSC in GenomicSEM*, n.d.). However, they have also indicated that progress on this development is limited by the paucity of multi-ancestry cohorts.

## Conclusions

This study demonstrates the viability of combining distinct EF tests from differing samples using a latent genetic factor. This will allow for larger meta-analytic GWAS of EF to be conducted in GenomicSEM without the need to unify EF measures across cohorts. However, while this study supports the existence of the unity-diversity structure of EF on the genetic level, the statistical power needed to fully model it for latent factor GWAS still poses a significant challenge at the current stage. Furthermore, our results suggest a need to reexamine the tests used to measure EF, particularly with respect to inhibition. The failure of the Stop-Signal to correlate genetically with other EF tests or load significantly onto latent EF factors limited the models that could be constructed, highlighting a need for greater theoretical clarity as to the relationship between EF tests and the latent construct. GWAS results suggest a potential relationship between various components of EF and psychiatric diagnoses. Furthermore, some results are suggestive of unique associations with specific EF domains, although additional research is needed to clarify these effects. In total, these results suggest that while single-test measures of EF may not be ideal in most circumstances, there is nonetheless value in conducting GWAS of such tests for their contribution to larger analyses of latent genetic factors.

## Acknowledgements and Data Availability

Special thanks to Emma Meaburn and Georgina Donati for providing the summary statistics for the ALSPAC GWAS, and Joel Gelernter and Huiping Zhang for providing the summary statistics for the Wisconsin Card Sorting Test.

Summary statistics will be made available through GWAS Catalog upon publication. FUMA results will likewise be made publicly available on that platform. Analysis code is available on Github: https://github.com/LCPerry/EFGSEM

