## Supplementary material for "GenomicSEM Modelling of Diverse Executive Function GWAS Improves Gene Discovery": Suplimental Methods

**Supplementary Methods**

Initially, our model included summary statistics from the Stop Signal reaction time, hypothesizing that its nonsignificant relationships with others in LDSC were just due to the GWAS's relatively low sample size and heritability producing large standard errors. However, we found that term behaved counterintuitively when included in the model. In all tested three-factor formulations, Stop Signal loaded weakly (~.10) and non-significantly onto inhibition. Similarly, in a bifactor formulation, Stop Signal loading onto common EF drops to nearly zero. While bifactor models of EF typically equate inhibition with common EF, the terms available to us did allow us to test models with inhibition specific variance. The version of that model where the ALSPAC working memory PC is allowed to cross load with inhibition (Supplemental Figure 1) achieves good fit by CFI and acceptable fit by SRMR, fitting better than the bifactor without an inhibition-specific factor. However, the loadings prove problematic. Stop Signal loads non-significantly and near zero to common EF, and while it dominates the IC factor, is loads onto it negatively, in the opposite direction of Stroop. Prospective Memory and the WM Principal component also load negatively, but both Stroop and Prospective memory are nonsignificant in their loadings. But dropping these two terms results in the remaining terms no longer loading significantly onto inhibition, and SRMR falls below acceptable fit.

The negative loadings onto inhibition from Stop Signal, ALSPAC working memory and prospective memory suggested that these three terms might be tapping a different factor. As such, we tested models that separated these three terms from Stroop onto a new factor, which we tentatively identify as response inhibition. Prospective memory also retained a cross loading onto the original inhibition with Stroop, as it could be conceptually compared to both Stop Signal and other inhibition tasks like Go/No-Go. The resulting four factor model (Supplemental Figure 2) tentatively supported this hypothesis because it showed significant and positive loadings for all terms, but fit statistics remained below acceptable thresholds (note that this model required the residual variance for the CHARGE DSST to be fixed to 0 to avoid empirical under-identification, as no EF terms correlated with the substitution factor). The bifactor variation was a dramatic improvement in fit, but reduced stop signal to the only significant loading onto either form of inhibition, rendering both inhibition factors empirically under-identified and leading us to reject the model. And while this could be considered consistent with a lack of inhibition-specific variance in bifactor models of EF, dropping the original inhibition and retaining only response inhibition causes the model to fall back below acceptable fit thresholds.

Ultimately, no tested models including Stop Signal produced both significant loadings for all terms and acceptable fit statistics. This suggested that the nonsignificant relationships between it and other EF measures in LDSC were due to more than just sample power, and that it may be tapping a construct separate from EF. As such, Stop Signal was dropped from the model. However, this analysis additionally suggested that prospective memory and ALSPAC working memory also tapped this construct on some level. Consequentially, we chose to account for it in the main model as a correlated residual between these two terms, which improved model fit.

Supplemental Table 1: Confirmatory factor analysis model results for the genomic structure of EF, with stop signal included

| Model | **$\chi^2$** | **df** | **$\chi^2$ p-value** | **AIC** | **CFI** | **SRMR** |
|---|---|---|---|---|---|---|
| Three Factor | 270.6759 | 32 | 1.345381e-39 | 316.6759 | 0.8874039 | 0.1628876 |

| | | | | | | |
|---|---|---|---|---|---|---|
| Three Factor + | 279.2928 | 31 | 9.498428e-42 | 327.2928 | 0.8828671 | 0.164469 |
| Three Factor with Substitution | 212.7092 | 28 | 2.632274e-30 | 266.7092 | 0.9128629 | 0.1563953 |
| Bifactor | 82.98456 | 28 | 2.378828e-07 | 136.9846 | 0.9740609 | 0.1464483 |
| Bifactor with Inhibition | 198.149 | 28 | 1.534849e-27 | 252.149 | 0.9197317 | 0.1550274 |
| Bifactor with Inhibition + | 63.03898 | 23 | 1.373198e-05 | 127.039 | 0.9811115 | 0.0992053 |
| Bifactor drop nonsig loadings from inhibition | 66.87524 | 25 | 1.108257e-05 | 126.8752 | 0.9802452 | 0.1138366 |
| Four Factor | 201.2959 | 23 | 1.951511e-30 | 265.2959 | 0.9158884 | 0.104868 |
| Bifactor two Inhibitions | 55.94492 | 19 | 1.663892e-05 | 127.9449 | 0.9825711 | 0.07772524 |
| Bifactor only Response Inhibition | 67.79836 | 24 | 4.688129e-06 | 129.7984 | 0.979338 | 0.1121269 |

+ denotes models with the ALSPAC WM Principal component cross loading onto both inhibition and working memory

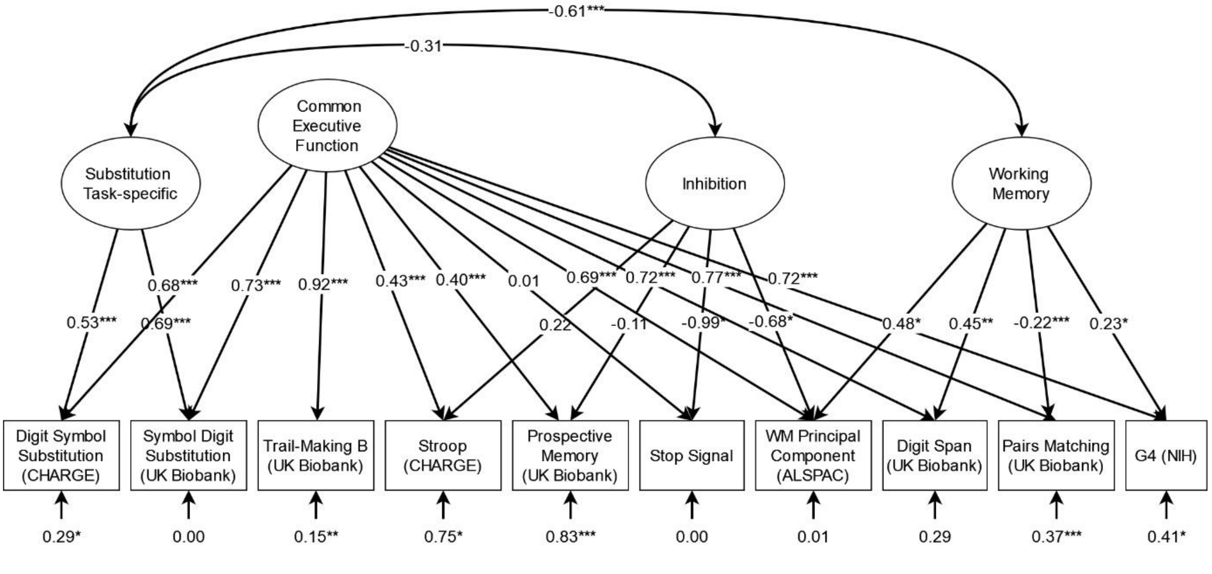


Supplemental Figure 1: Standardized results from the bifactor model including stop-signal and an inhibition-specific factor. *p < .05, ** p < .01, *** p < .001

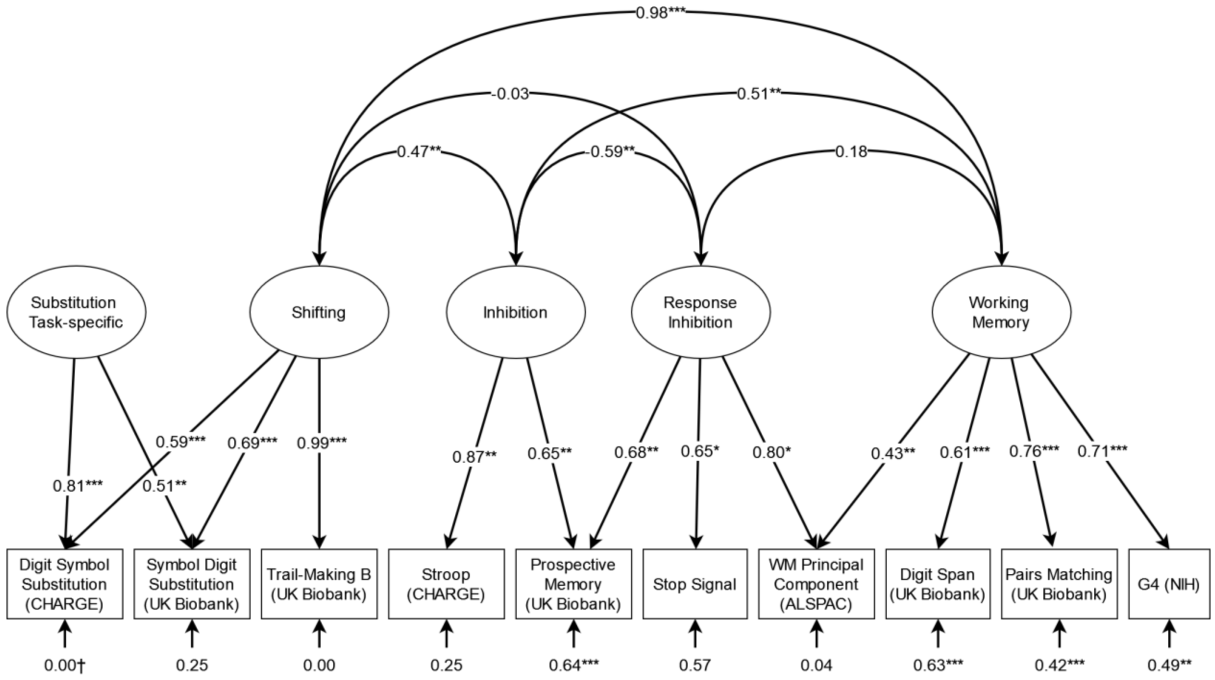


Supplemental Figure 2: Standardized results from the correlated factors model including the response inhibition factor. Correlations between the substitution-specific and inhibition, response inhibition and working memory factor are present but non-significant in the model, and dropped here for ease of visualization. † indicates that residual variance was constrained to 0, to prevent empirical under-identification. *p < .05, ** p < .01, *** p < .001
